# AIBP controls TLR4 inflammarafts and mitochondrial dysfunction in a mouse model of Alzheimer’s disease

**DOI:** 10.1101/2024.02.16.580751

**Authors:** Yi Sak Kim, Soo-Ho Choi, Keun-Young Kim, Juliana M. Navia-Pelaez, Guy A. Perkins, Seunghwan Choi, Jungsu Kim, Nicolaus Nazarenkov, Robert A. Rissman, Won-Kyu Ju, Mark H. Ellisman, Yury I. Miller

## Abstract

Microglia-driven neuroinflammation plays an important role in the development of Alzheimer’s disease (AD). Microglia activation is accompanied by the formation and chronic maintenance of TLR4 inflammarafts, defined as enlarged and cholesterol-rich lipid rafts serving as an assembly platform for TLR4 dimers and complexes of other inflammatory receptors. The secreted apoA-I binding protein (APOA1BP or AIBP) binds TLR4 and selectively targets cholesterol depletion machinery to TLR4 inflammaraft expressing inflammatory, but not homeostatic microglia. Here we demonstrated that amyloid-beta (Aβ) induced formation of TLR4 inflammarafts in microglia in vitro and in the brain of APP/PS1 mice. Mitochondria in Apoa1bp^-/-^ APP/PS1 microglia were hyperbranched and cupped, which was accompanied by increased ROS and the dilated ER. The size and number of Aβ plaques and neuronal cell death were significantly increased, and the animal survival was decreased in Apoa1bp^-/-^ APP/PS1 compared to APP/PS1 female mice. These results suggest that AIBP exerts control of TLR4 inflammarafts and mitochondrial dynamics in microglia and plays a protective role in AD associated oxidative stress and neurodegeneration.

## INTRODUCTION

Neuroinflammation and activated microglia, as well as alterations in cholesterol metabolism play important roles in the development of Alzheimer’s disease (AD). Cholesterol and many receptors governing inflammatory responses colocalize in the ordered membrane microdomains, often designated as lipid rafts. Lipid rafts are essential for cell physiology as they provide a solid, cholesterol-enriched platform within a membrane, where molecular complexes, such as toll-like receptor-4 (TLR4) dimers, can assemble without battling forces of chaos in the disorderly liquid phase of the surrounding membrane^1^. Lipid rafts in quiescent cells are highly dynamic and transient^2^. Upon cell activation, lipid rafts cluster and become more stable to accommodate agonist-induced assembly of receptors into functional complexes and the initiation of signaling^3^. For example, in LPS-stimulated microglia, TLR4 dimer-hosting lipid rafts last for 15 min and then diminish due to internalization of the LPS-TLR4 complex^4^. However, under chronic inflammatory conditions, TLR4 dimers and increased lipid raft levels are surprisingly persistent; they can be found in spinal microglia 21 days after a chemotherapeutic intervention, which results in peripheral neuropathy and a persistent pain state^5^. Treatments that reduce microglial lipid rafts alleviate tactile allodynia, whereas increasing the membrane levels of cholesterol and lipid rafts promotes allodynia^5^. Enlarged lipid rafts, TLR4 dimers, and assemblies of other inflammatory receptors persist also in macrophages isolated from mouse atherosclerotic lesions, which is another example of a chronically inflamed tissue^6^.

To conceptualize the findings of persistent lipid rafts under chronic inflammatory conditions, we introduced the term inflammarafts, defined as enlarged/clustered lipid rafts harboring activated receptors and adaptor molecules and serving as a scaffold to organize cellular inflammatory response^7^. The TLR4 inflammaraft expression is associated with hyperinflammatory microglia/macrophage reprogramming^4–6^. Furthermore, one agonist can prime macrophages – via formation of inflammarafts that host a community of different inflammatory receptors – for a hyperinflammatory response to a spectrum of secondary stimuli^6^. The AD brain is flooded with amyloid-beta (Aβ) and other disease-associated molecular patterns (DAMPs) that provide fertile ground for maintaining inflammarafts in microglia reprogrammed for a low-grade but persistent expression of inflammatory genes and reactive oxygen species (ROS).

ApoA-I binding protein (AIBP; encoded by the *APOA1BP* gene, a.k.a. *NAXE*) is a 32 kDa protein, which depending on phosphorylation by PKA, can be intracellular or secreted^8, 9^. In humans, eQTL rs4661188 for *APOA1BP* showed significant inverse interaction with family history of AD^10^, and *APOA1BP* has been identified as a susceptibility locus for migraine^11^. AIBP is an important regulator of cellular cholesterol metabolism, which facilitates cholesterol depletion from the plasma membrane and selective disruption of inflammarafts^5, 12^. The AIBP selectivity toward TLR4 inflammarafts is due to its binding to TLR4 and thus directing cholesterol depletion to the cells expressing high surface levels of TLR4, such as activated microglia and nociceptive dorsal root ganglia neurons^4, 13^. Intrathecal administration of the recombinant AIBP protein reduces TLR4 dimerization and lipid raft levels, the key characteristics of TLR4 inflammarafts, and alleviates tactile allodynia in a mouse model of chemotherapy-induced peripheral neuropathy^5^.

In the present study we demonstrate that in an APP/PS1 model of AD, AIBP deficiency significantly exacerbates microglial TLR4 inflammarafts and mitochondrial dysfunction, oxidative stress, Aβ plaque accumulation and neuronal cell death. These results reveal a new mechanism in which the TLR4 inflammaraft – mitochondrial dysfunction axis in microglia propagates oxidative stress, and suggest a neuroprotective function of AIBP in the AD brain.

## RESULTS

### Amyloid beta induces TLR4 inflammarafts expression and oxidative stress in microglia

We previously demonstrated that exposure to LPS, a TLR4 agonist, or to Pam3CSK4, a TLR2 agonist, induces TLR4 inflammarafts in macrophages and microglia^4–6^. We asked whether Aβ can similarly induce TLR4 inflammarafts. BV-2 microglia were analyzed after a 24-hour incubation with Aβ oligomers or LPS. Both stimuli induced robust increases in the levels of lipid rafts and surface expression of TLR4, which localized to lipid rafts (**Figures 1A and 1B**), without affecting cell viability (**Figure S1A**). To further characterize TLR4 inflammarafts, we conducted a proximity ligation assay (PLA) with antibodies against TLR4 and cholera toxin B (CTxB, which binds to ganglioside GM1, a lipid raft component), detecting their juxtaposition at a distance of <40nm. In BV-2 microglia, Aβ increased TLR4 localization to lipid rafts (TLR4-LR) as measured by PLA (**Figures 1C, 1D and S1B**). Using a flow cytometry assay^4–6^, we detected increased levels of TLR4 dimers in BV-2 microglia exposed to Aβ, as well as increased expression of TLR4 and lipid rafts (**Figure 1E**). Importantly, in parallel with the Aβ-induced TLR4 inflammarafts, BV-2 microglia also displayed high levels of ROS in response to Aβ (**Figure 1F**).

**Figure 1.**
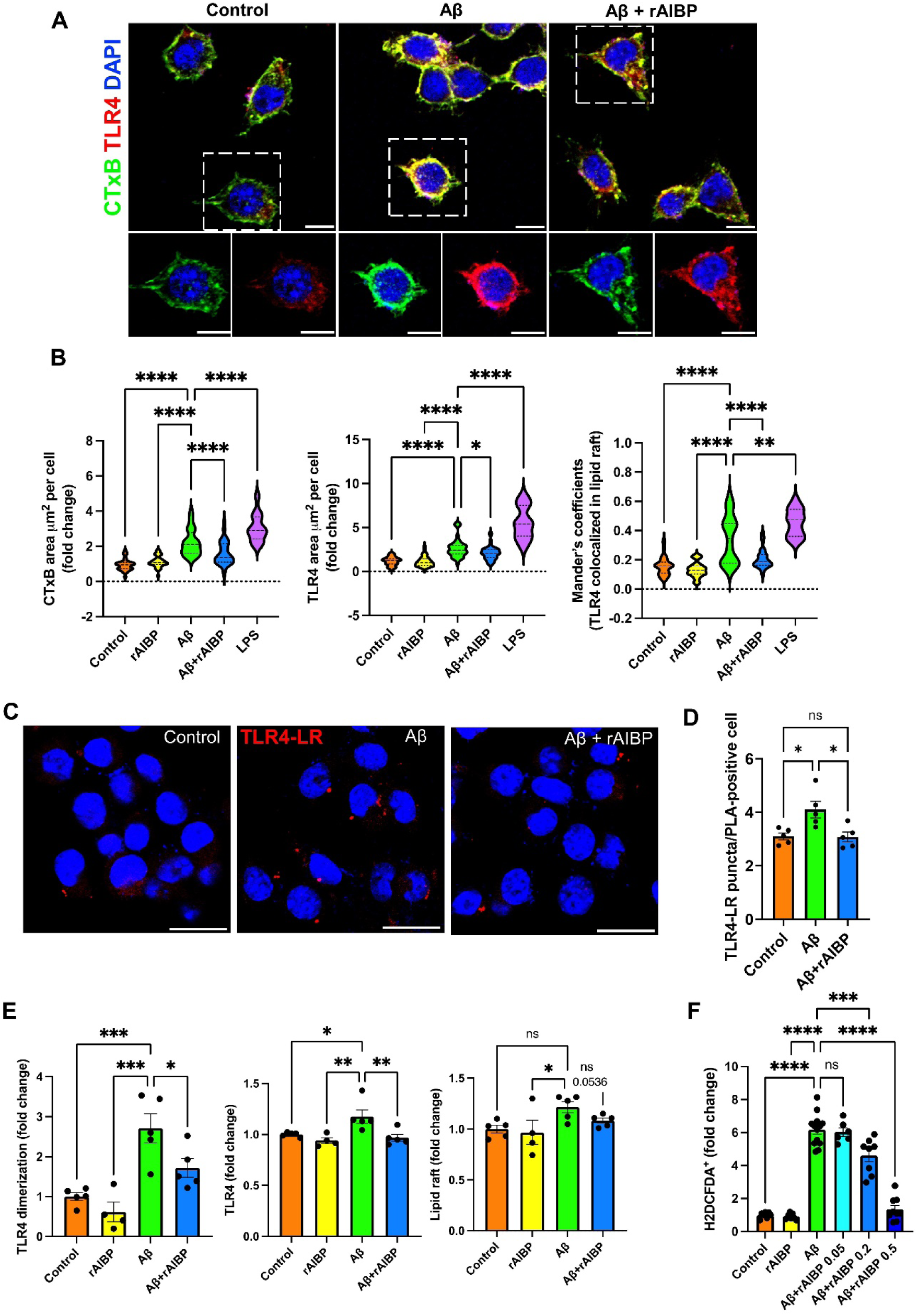
AIBP protects microglia against Aβ-induced TLR4 inflammarafts and oxidative stress. BV-2 cells were incubated with rAIBP for 2 hours before a 24-h exposure to Aβ (7PA2-conditioned media containing 300 pM Aβ_42_) or control (CHO-conditioned media). The rAIBP concentrations were 0.5 µg/ml in panels A-E, and 0.05, 0.2, or 0.5 µg/ml in panel F. See also **Figure S1A**. (**A** and **B**) Colocalization of TLR4 and lipid rafts. Immunostaining with CTxB-FITC (lipid raft; green), anti-TLR4 antibody (red), and DAPI (nuclei; blue). As a positive control, BV-2 cells were incubated with 100 ng/ml LPS. Manders’ overlap coefficients were calculated to assess TLR4 overlapping with lipid rafts. Data were collected from 24-43 fields of 4-8 biological replicates in three independent experiments. Scale bar, 10 μm, (**C** and **D**) TLR4-CTxB (LR, lipid rafts) proximity ligation assay (Texas red probe) in BV-2 cells. Scale bars, 20 μm. The number of PLA puncta signals in PLA-positive cells was quantified from an average of two fields in each biological replicate (*n* = 5/group), from two independent experiments. See also **Figure S1B**. (E) Flow cytometry analysis of TLR4 expression and dimerization using TLR4-APC and TLR4/MD2-PE antibodies and of lipid raft content (CTxB-FITC) in BV-2 cells; 4-5 biological replicates per group from two independent experiments. (F) Flow cytometry analysis of intracellular ROS (H2DCFDA); 7-13 biological replicates per group, from three independent experiments. Mean±SEM. One-way ANOVA with Dunnett’s multiple comparisons test (Aβ vs. each group).

AIBP selectively targets TLR4 inflammarafts via binding to TLR4 and enhancing cholesterol depletion from the plasma membrane^4, 5^. Adding recombinant AIBP protein (rAIBP) to BV-2 microglia significantly inhibited Aβ-induced increases in lipid rafts, TLR4 expression and localization to lipid rafts, TLR4 dimerization and generation of ROS (**Figures 1A-1F**). These results suggest a protective role of AIBP under conditions of Aβ-induced inflammation and oxidative stress.

### AIBP deficiency exacerbates TLR4 inflammarafts and oxidative stress in vivo

To test the proposed protective role of AIBP in vivo, we crossed *Apoa1bp^-/-^* mice^14, 15^ with APP/PS1 transgenic mice [expressing the mutant human APP751 (KM670/671NL, Swedish) and the mutant human PSEN1 (M146L)]^16^, and compared them with APP/PS1 and wild type (WT) mice. The AIBP deficiency in *Apoa1bp^-/-^* APP/PS1 mice was confirmed by RT-qPCR (**Figure S1C**). At 6 months of age, CD11b^+^CD45^low^ brain microglia from APP/PS1 mice displayed a trend toward increases in TLR4 dimers and the levels of lipid rafts and total TLR4. This trend was further amplified in the microglia from *Apoa1bp^-/-^* APP/PS1 mice, resulting in significant differences in TLR4 expression and dimerization, compared to WT mice (**Figures 2A-2C**). In addition, AIBP deficiency significantly exacerbated oxidative stress in the APP/PS1 mouse brain, as detected with DHE in both hippocampus and cortex regions (**Figures 2D and 2E**).

**Figure 2.**
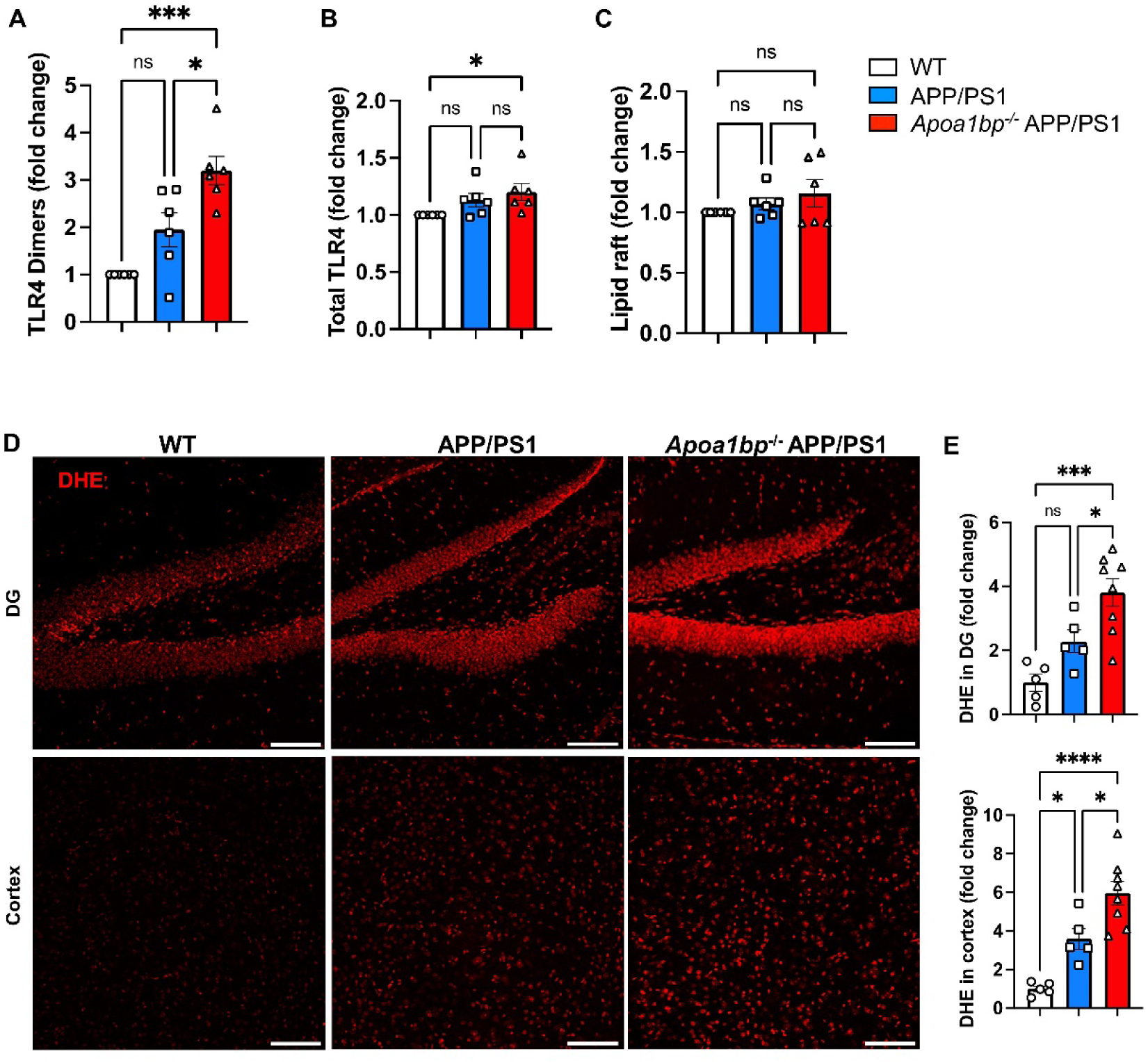
AIBP deficiency exacerbates TLR4 inflammarafts and oxidative stress in APP/PS1 mice. (**A**-**C**) Flow cytometry analysis of CD11b^+^ CD45^low^ microglia cells in single cell suspensions isolated from age-matched 6-months old female mice (*n*=6 per group). (**A**) TLR4 dimerization, (**B**) TLR4 expression, and (**C**) lipid raft content (CTxB). (**D** and **E**) Female mouse brain tissues (WT *n*=5, APP/PS1 *n*=5, and *Apoa1bp*^-/-^ APP/PS1 *n*=8) were stained with the superoxide indicator dihydroethidium (DHE, 10 μM; red). Representative images in the dentate gyrus (DG) and cortex. Scale bar, 100 μm. Mean±SEM. One-way ANOVA with Tukey’s multiple comparisons test. See also **Figure S1C**.

### Distorted mitochondrial and ER architecture in *Apoa1bp^-/-^* APP/PS1 microglia

Evidence of exacerbated oxidative stress suggests a possibility of mitochondrial dysfunction in *Apoa1bp^-/-^* APP/PS1 microglia. Using a serial blockface scanning electron microscopy (SBEM) protocol for imaging hippocampal microglia, we found smaller and branched mitochondria in APP/PS1 compared to WT mice (**Figures 3A-3J**). Remarkably, *Apoa1bp^-/-^* APP/PS1 mitochondria were hyper-branched and large because they had a greater number of branches per mitochondrion and were more cupped, the cupping shape itself adding to the volume (**Figures 3A-3J**). The increased volume of *Apoa1bp^-/-^* APP/PS1 compared to APP/PS1 mitochondria may suggest an increase in energy demand^17^ in the absence of AIBP. However, the distorted morphology, and specifically the cup-shaped mitochondria point to an increased generation of ROS^18–23^. Ring- and cup-shaped mitochondria were found in human AD brain microglia as well (**Figures 3K and 3L**), suggesting the relevance of the findings of cupped mitochondria in *Apoa1bp^-/-^* APP/PS1 microglia to human AD pathology.

**Figure 3.**
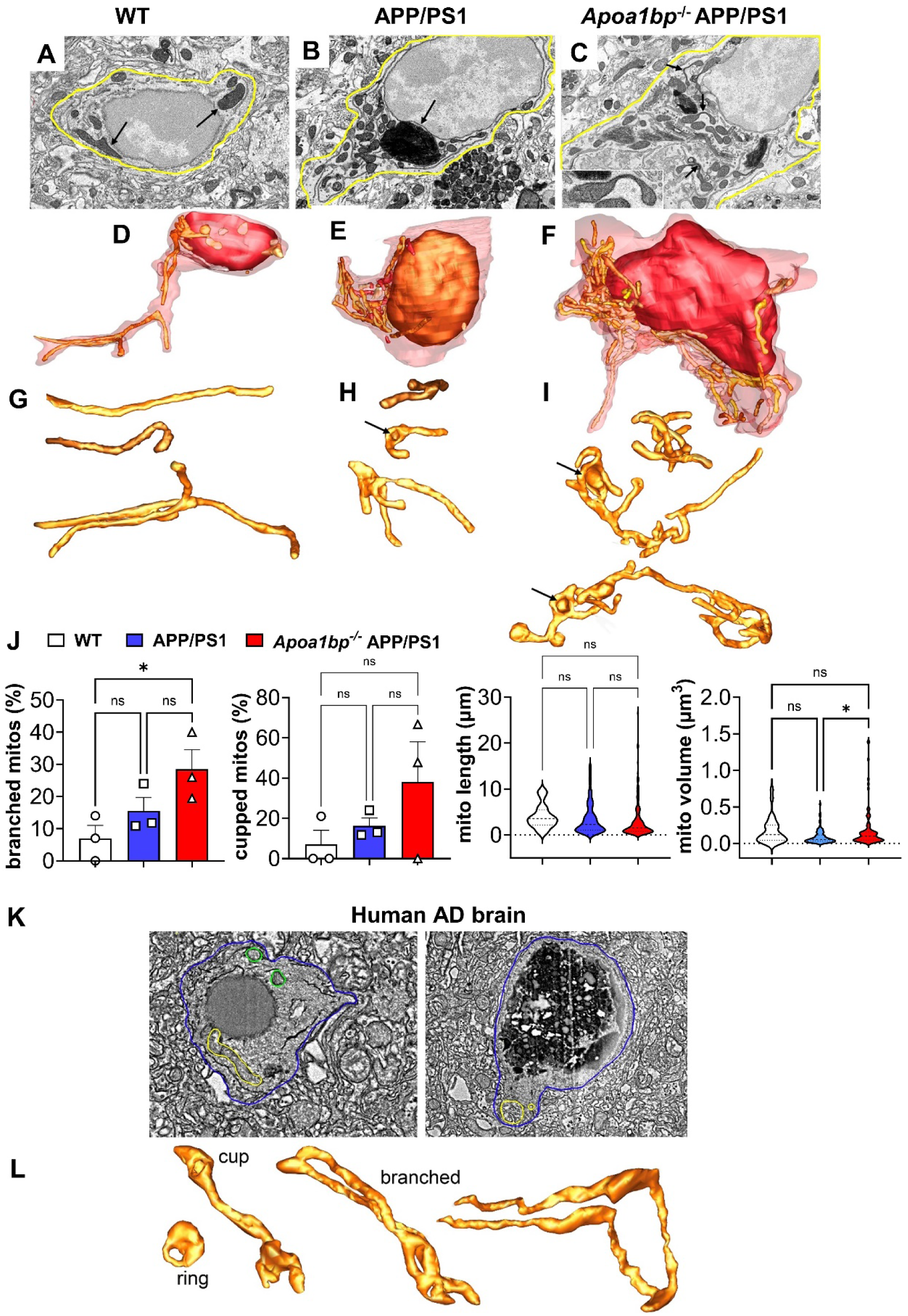
Hyper-branched and cupped mitochondria in microglia of *Apoa1bp^-/-^* APP/PS1 mice and AD subjects. SBEM slices and surface rendering of mitochondria from hippocampal microglia of WT (**A, D, G**), APP/PS1 (**B, E, H**) and *Apoa1bp^-/-^* APP/PS1 (**C,F, I**) female mice. (**A-C**) SBEM slices; arrows in **A**: large mitochondria; **B**: small mitochondria near a lipofuscin aggregate; **C** and inset: cupped-shaped mitochondria. (**D-F**) Segmentation and surface-rendering of the microglia cell shown in **A-C**, displaying the cell volume boundary (translucent maroon), the nucleus (brown) and mitochondria (lighter shades of brown). Fourteen mitochondria in **D**, 46 in **E** and 72 in **F**. Higher mitochondrial volume density in *Apoa1bp^-/-^* APP/PS1 than in either WT or APP/PS1 microglia. (**G-I**). Mitochondria from the surface-rendering shown in **D-F**. (**G**) Typically large mitochondria in WT; a few are branched. (**H**) APP/PS1: smaller mitochondria, with more than twice as many branched mitochondria, than typically found in WT microglia, but those branched have fewer branches, contributing to their smaller volumes. A greater number of cupped mitochondria (arrow). (**I**) *Apoa1bp^-/-^* APP/PS1: increased branching and cupping (arrows) compared with WT and APP/PS1 microglia, which contribute to the large mitochondrial volumes. (J) Measurements of branched and cupped mitochondria, and total length and volume (n=3-6 microglia from 2 mice per group; a total of 28 (WT), 102 (APP/PS1) and 166 (*Apoa1bp^-/-^* APP/PS1) mitochondria were measured). (K) Human AD cortex SBEM showing 2 microglia with 2 branched mitochondria on the left (yellow and green traces) and the ring-shaped mitochondrion on the right (yellow trace). The cell membrane is traced in blue. (L) Surface rendering of ring, cupped and branched mitochondria from SBEM volumes shown in panel **K**.

The results showing distorted mitochondrial architecture in brain microglia under conditions of increased Aβ production in APP/PS1 mice, which was further exacerbated by AIBP deficiency (**Figures 3A-3J**), suggest that AIBP plays a protective role in mitochondrial respiration. To test this hypothesis, we measured oxygen consumption rate (OCR) in BV-2 microglia exposed to Aβ, in the absence and presence of rAIBP. Basal and maximal respiration, spare capacity and ATP production were all significantly decreased in Aβ challenged microglia. However, in the presence of rAIBP, all these parameters in Aβ challenged microglia were indistinguishable from those in control, unchallenged cells (**Figure 4**).

**Figure 4.**
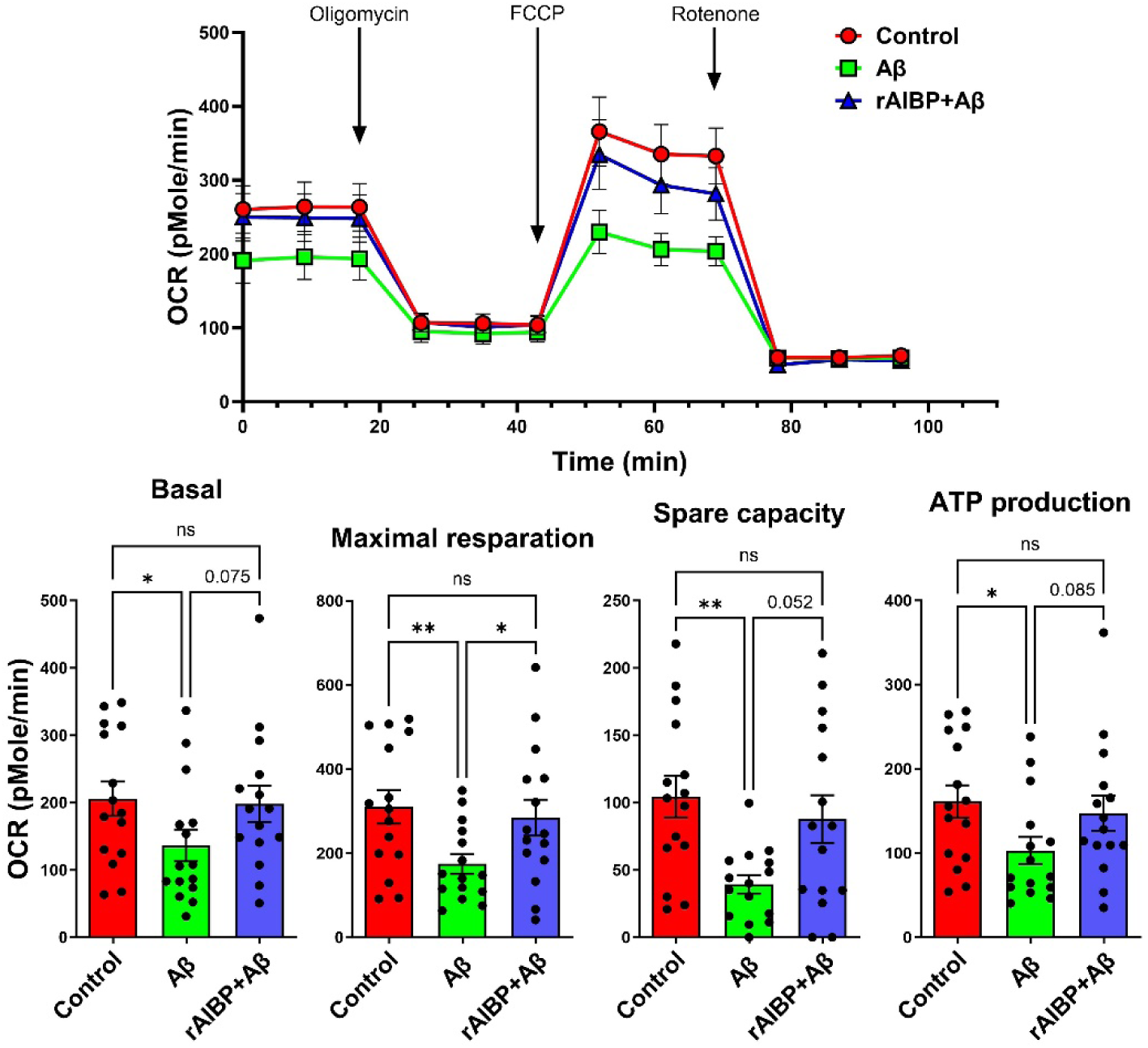
AIBP protects microglia against Aβ-altered mitochondrial respiration. BV-2 cells were preincubated with rAIBP (0.2µg/ml) or BSA for 2 h before a 24-h exposure to Aβ (7PA2-conditioned media containing 300 pM Aβ_42_) or control (CHO-conditioned media). Seahorse data are OCR plotted versus time. Basal and maximal respiration, spare capacity, and ATP production were measured (*n=*15 per group). Mean±SEM. One-way ANOVA with Tukey’s multiple comparisons test.

In addition to mitochondrial architecture, we examined the endoplasmic reticulum (ER) in hippocampal microglia from APP/PS1 and *Apoa1bp^-/-^* APP/PS1 mice. Activated microglia phenotype was evident in the proximity of dystrophic neurites; however, there was no evidence of the dilation in ER cisternae from APP/PS1 compared to WT microglia. In contrast, the ER cisternae in *Apoa1bp^-/-^* APP/PS1 microglia showed a swollen morphology compared to the WT and APP/PS1 microglia, with the ER dilation particularly profound near the amyloid plague (**Figure 5**).

**Figure 5.**
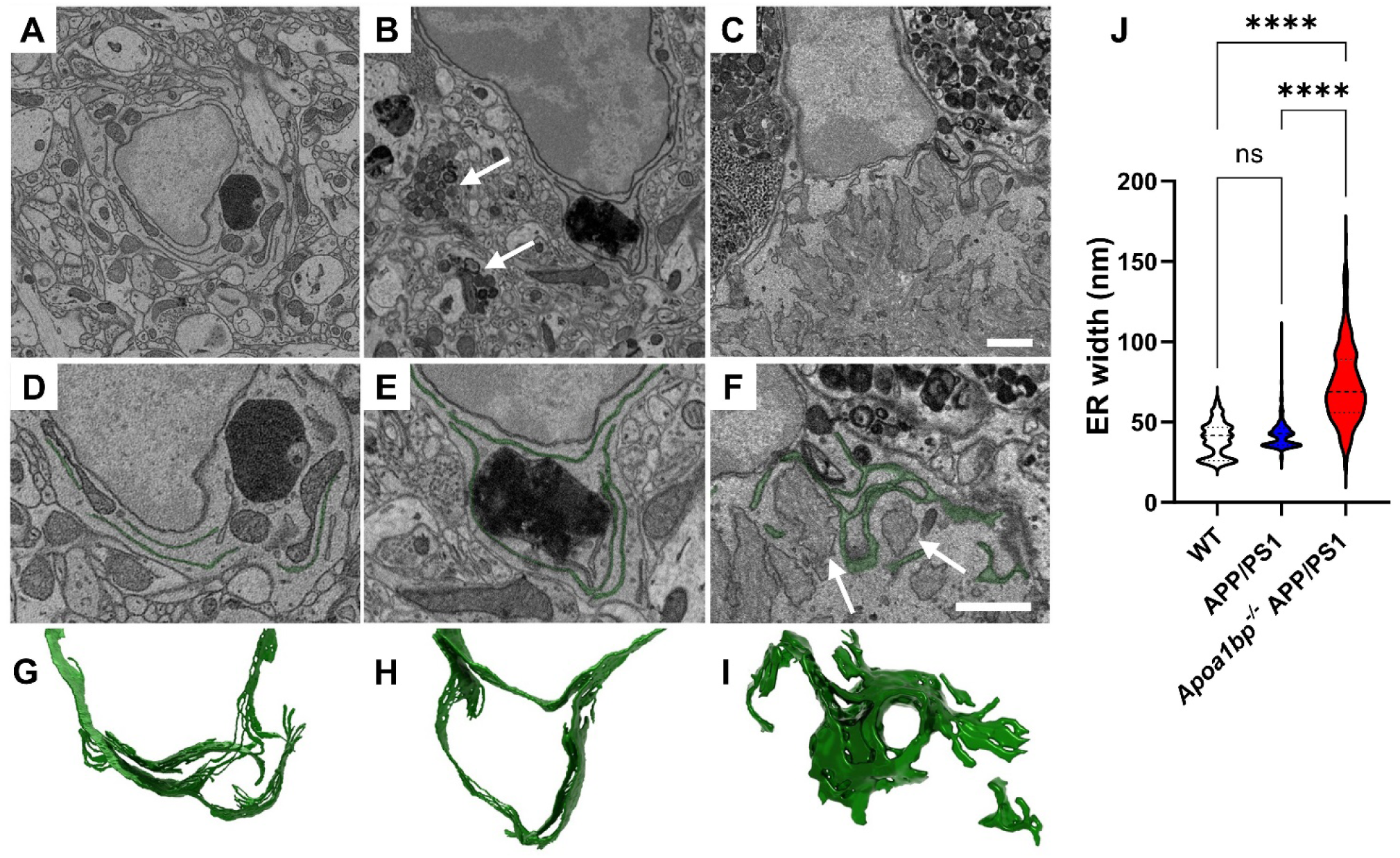
AIBP deficiency triggers ER swelling in activated microglia. (**A, D, G**) A representative SBEM image plane from female WT microglial cell showing a typical ER structure (**A**). Images for cross-section (**D**) and 3D rendering (**G**) of thin ER cisternae in WT microglia. (**B, E, H**) Activated microglia phenotype and dystrophic neurites (arrows) in female APP/PS1 mice, showing no evidence of the dilation in ER cisternae. (**C, F, I**) Swollen ER cisternae morphology in microglia from female *Apoa1bp^-/-^* APP/PS1 mice. Note that ER dilation was profound near the amyloid plague (arrows). (**J**) Quantification of the ER width in female WT, APP/PS1, and *Apoa1bp^-/-^* APP/PS1 microglia; violin plot (*n*=250-259 per group; lumen width was measured for 68-96 ER segments per microglial cell; 3 microglial cells from 2 mouse brains per group). Mean±SEM. One-way ANOVA with Tukey’s multiple comparisons test. Scale bar, 1µm.

### Exacerbated neuropathology in AIBP deficient APP/PS1 mice

The in vitro experiments showing protective role of rAIBP (**Figures 1 and 4**) and the increased TLR4 inflammarafts, distorted microglial mitochondria, ER stress, and oxidative stress in *Apoa1bp^-/-^* APP/PS1 brain (**Figures 2**, **3 and 5**) suggest that AIBP deficiency might result in exacerbated AD-like pathology. Unchallenged *Apoa1bp^-/-^* mice are viable, with a normal life span and no neuroanatomical or behavioral abnormalities^5, 14, 15, 24^. There was no microgliosis, changes in microglia morphology, nor Aβ deposition in the brain of *Apoa1bp^-/-^* mice (**Figures 6A**, **6B**, **7A**, **7B and S2A**). However, *Apoa1bp^-/-^* APP/PS1 mice at 6 months of age had microgliosis and the microglia characterized by significantly fewer and shorter branches and larger soma, compared with the microglia in WT, *Apoa1bp^-/-^* or APP/PS1 brain (**Figure 6**). In addition, there was a significantly larger number and total area of Aβ plaques in the hippocampus of *Apoa1bp^-/-^* APP/PS1 compared to APP/PS1 mice; the difference was particularly significant in female mice (**Figures 7A, 7B and S2A**). There were also more microglia associated with Aβ plaques in *Apoa1bp^-/-^* APP/PS1 female mice (**Figure S2B**). Cortex of *Apoa1bp^-/-^* APP/PS1 female mice contained more insoluble Aβ_42_ peptides compared to that in APP/PS1 mice (**Figure 7C**). The activated microglia phenotype and an increased number of Aβ plaques were associated with significantly increased neuronal cell death in *Apoa1bp^-/-^* APP/PS1 compared to APP/PS1 mice (**Figures 8A**, 8B **and S3A-3C**). The accelerated neuronal cell death was associated with lower survival rates of *Apoa1bp^-/-^* APP/PS1 mice (**Figure 8C**).

**Figure 6.**
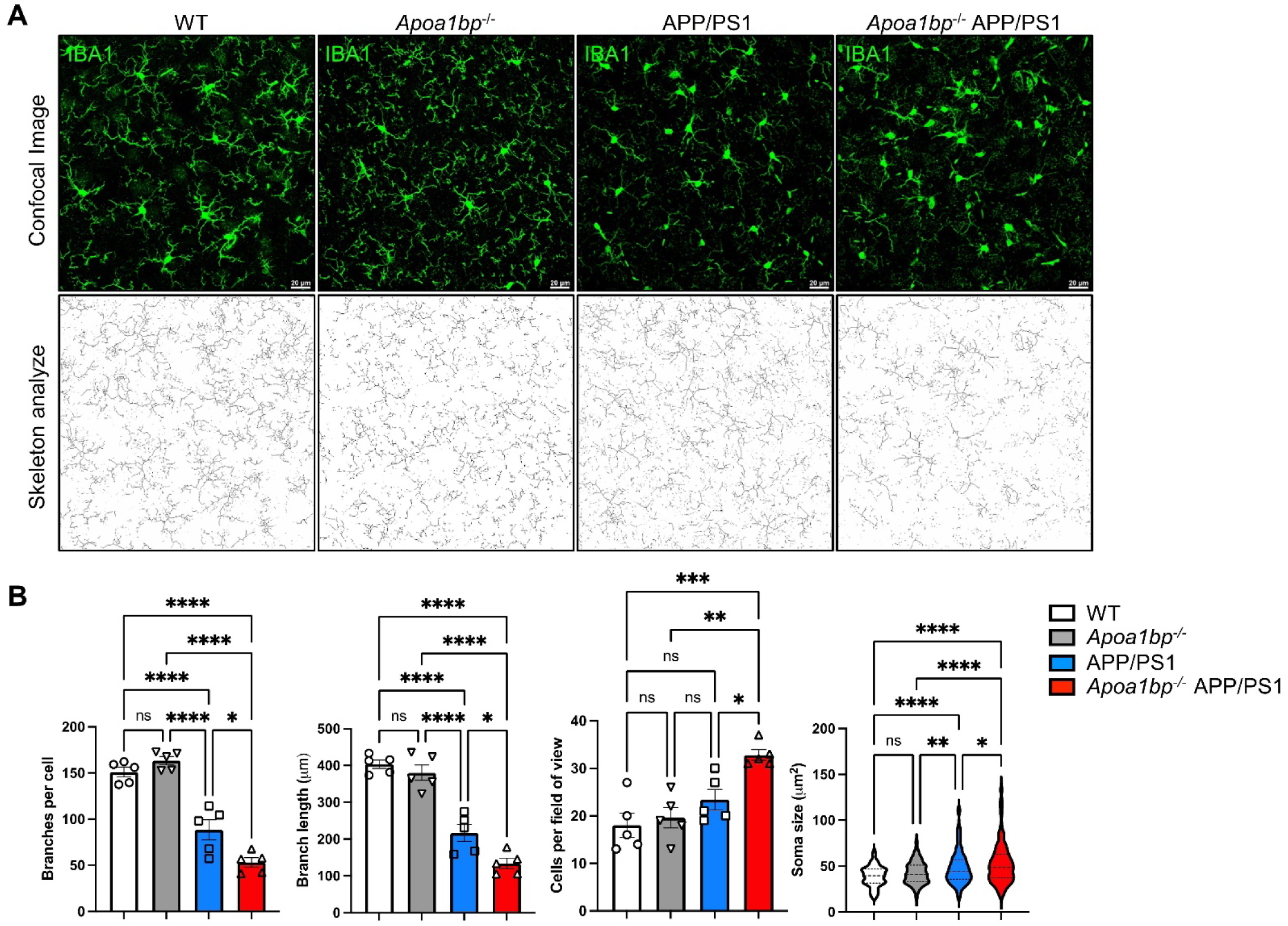
Altered microglia morphology in *Apoa1bp^-/-^* APP/PS1 mouse brain. (**A**) Representative IBA1-positive microglia cells (green) and skeletonized rendering in the cortex of 6 months- old female mice. Scale bar, 20 μm. (**B**) Skeletonize macros in ImageJ were used to measure the number of branches, branch length, the number of cells per field of view, and the soma size (195-313 microglial cells per group). An average of two fields of view for each mouse (*n*=5 per group). Mean±SEM. One-way ANOVA with Tukey’s multiple comparisons test.

**Figure 7.**
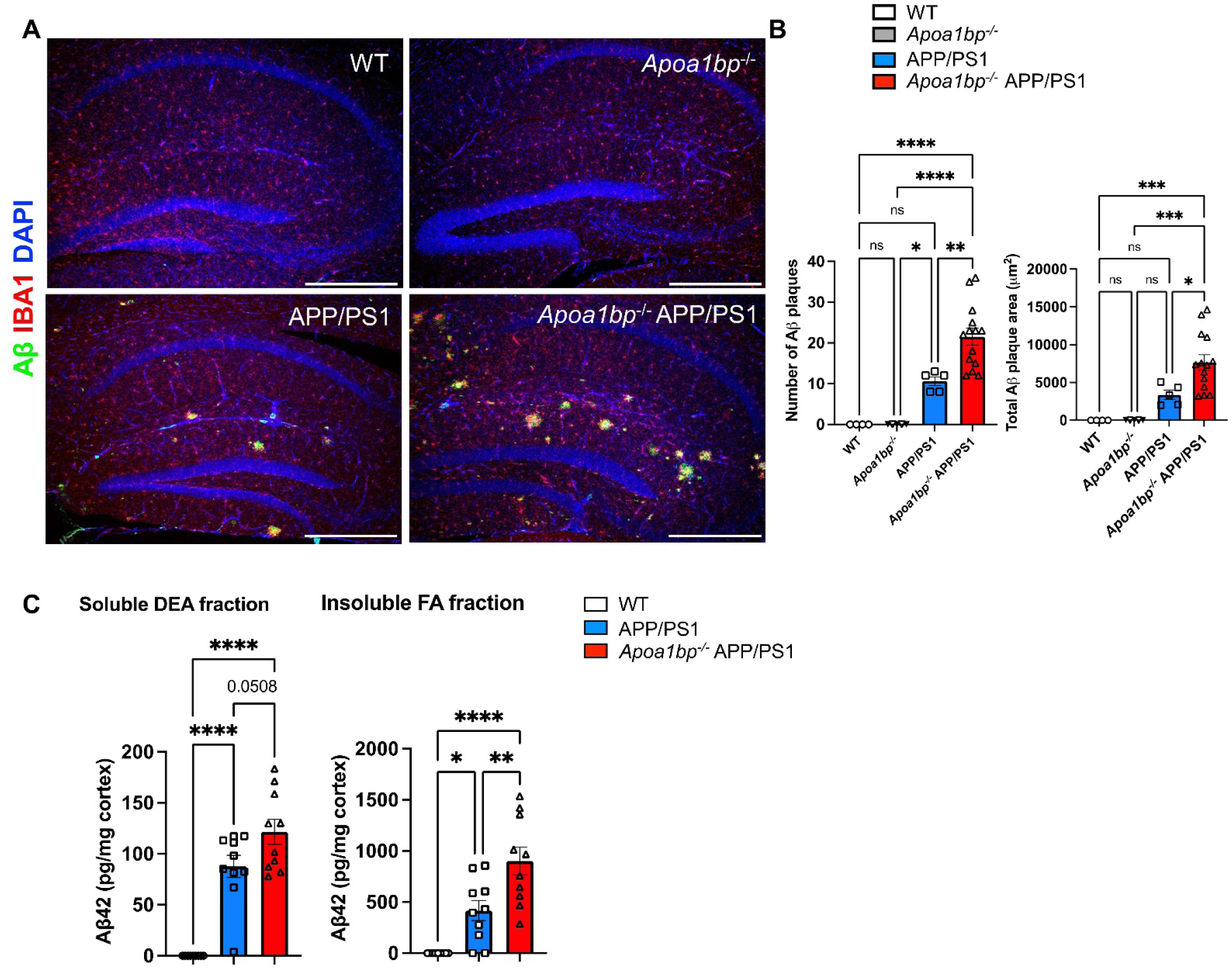
AIBP deficiency increases number of Aβ plaques in APP/PS1 mouse brain. (**A**) Representative hippocampal images showing 82E1 (Aβ; green), IBA1 (microglia; red), and DAPI (nuclei; blue) staining in the brain of age-matched 6 months-old female mice. Scale bar, 400 μm. (**B**) The number of Aβ plaques and the total plaque area in the hippocampus. WT (*n*=4), *Apoa1bp^-/-^* (*n*=5), APP/PS1 (*n*=5), and *Apoa1bp^-/-^* APP/PS1 (*n*=14). (**C**) Cortical Aβ_42_ peptide levels in age-matched 6 months-old female mice: WT (*n*=9), APP/PS1 (*n*=10), and *Apoa1bp^-/-^* APP/PS1 (*n*=10) measured in soluble DEA fraction and insoluble FA fraction. Mean±SEM; one-way ANOVA with Tukey’s multiple comparison test. See also **Figure S2**.

**Figure 8.**
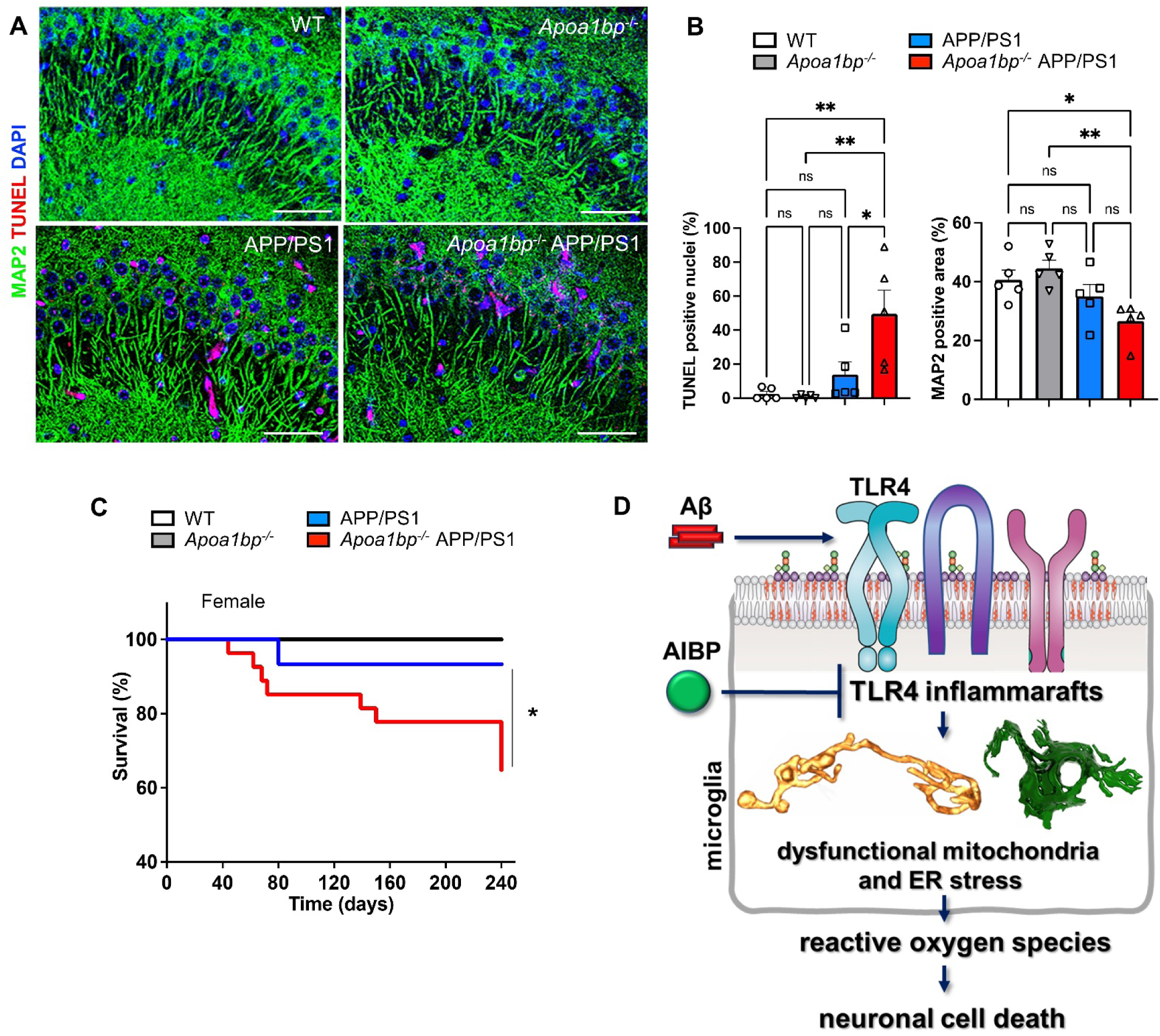
AIBP deficiency exacerbates neuronal cell death and reduces survival of APP/PS1 mice. (**A** and **B**) TUNEL (red) and MAP2 (green) staining in the CA3 area of age-matched 6 months-old female mice (*n*=5). A: Representative images. Scale bar, 50 μm. B: the percentage of TUNEL-positive nuclei and MAP2-positive area. See also **Figure S3**. (**C**) Survival curve of female mice: WT (*n*=10), *Apoa1bp^-/-^* (*n*=7), APP/PS1 (*n*=14), and *Apoa1bp^-/-^* APP/PS1 (*n*=20). Mean±SEM; one-way ANOVA with Tukey’s multiple comparison test (**B**) and Mantel-Cox log-rank test (**C**). (**D**) This study results suggest that increasing levels of Aβ induce TLR4 inflammaraft formation in microglia, which can be inhibited by adding recombinant AIBP protein in vitro, or exacerbated by AIBP deficiency in vivo. The Aβ overproduction and AIBP deficiency together lead to dysregulation of mitochondrial dynamics and the appearance of hyper-branched and cupped mitochondria, as well as dilated ER, both characteristic of the conditions associated with oxidative stress. Increased ROS production in the brain induces neuronal cell death.

## DISCUSSION

In this study, we applied a novel framework of assessing TLR4 inflammarafts, mitochondrial dysfunction and oxidative stress, to test the hypothesis that AIBP plays a protective role in Aβ-associated neuropathology. The premise for this work was the notion that persistent maintenance of TLR4 inflammarafts – enlarged lipid rafts hosting assemblies of TLR4 and other inflammatory receptors – in brain microglia is an important factor in chronification of neuroinflammation and oxidative stress, abetting Aβ plaque growth and neuronal cell death. We have demonstrated that, like LPS, Aβ induced formation of TLR4 inflammarafts in microglia, which was associated with mitochondrial dysfunction and oxidative stress. Adding rAIBP nullified these effects. Accordingly, in an Aβ mouse model (APP/PS1 mice), the AIBP deficiency exacerbated microglial inflammarafts, alteration in mitochondrial and ER architecture, and oxidative stress. These changes led to increased Aβ plaque formation and the increased neuronal cell death (**Figure 8D**) and accompanied reduced survival of *Apoa1bp^-/-^* APP/PS1 mice.

Aβ oligomers and other DAMPs engage inflammatory receptors located in lipid rafts and induce fusion of isolated lipid rafts into inflammarafts enriched with TLR4 dimers and likely other receptor complexes. These processes initiate cell reprogramming, including alterations in cholesterol and lipid metabolism, which help maintain high levels of cholesterol, sphingolipids and phospholipids with long saturated fatty acyl chains – all the components of lipid rafts^5, 7, 25^. The exact genetic, epigenetic and metabolic mechanisms of chronic TLR4 inflammaraft maintenance in microglia exposed to Aβ remain to be elucidated. However, the finding of increased TLR4 inflammarafts in AD microglia is consistent with the well characterized association of AD with dysregulation of cholesterol and lipid metabolism^26–30^.

Specifically, both TLR4 activation and dysregulation of cellular cholesterol metabolism result in mitochondrial morphological changes and dysfunction^29, 31, 32^. Accordingly, AIBP deficiency, which exacerbated TLR4 inflammarafts, was characterized by hyperbranched and cupped mitochondria. Instances of hyperbranched and cup-shaped mitochondria have been reported. For example, hyperbranched mitochondria in human urinary podocyte-like epithelial cells expressing genetic variants of the *APOL1* gene were accompanied by an increase in the mRNA level of the mitochondrial fusion protein MFN1^33^. The observation of cup-shaped mitochondria in brain microglia points to the possibility of an increased generation of reactive oxygen species and ensuing oxidative stress in *Apoa1bp^-/-^* APP/PS1 mice, leading to exacerbated pathology in AIBP-deficient tissues. Indeed, several studies, in senile dogs’ parathyroid^18^, Schwann cells^19^, alcohol, venom or chemically induced liver toxicity^20–22^, and carcinogen-induced renal oncogenesis^23^ made the connection between cup-shaped mitochondria and oxidative stress. Thus, we interpret the appearance of cup-shaped mitochondria in cells from AIBP deficient mice as the cause of oxidative stress, documented in the brain of *Apoa1bp^-/-^* APP/PS1 mice.

In addition, our SBEM studies demonstrated that microglial cell bodies in *Apoa1bp^-/-^* APP/PS1 mice were larger and displayed ultrastructural signs of cellular stress, particularly near amyloid plaques. ER stress is a prominent indicator of cellular stress at the subcellular level, which is read out at high spatial resolution as a lumen dilation of ER cisternae. Dilated ER has been reported in several neurodegenerative diseases, including amyotrophic lateral sclerosis and AD^34^. Our findings indicate a significant increase in the ER lumen expansion within microglial cell bodies of *Apoa1bp^-/-^* APP/PS1 mice compared to APP/PS1 or WT mice. These findings suggest that AIBP deficiency triggers ER stress, which is a causative factor in the development of AD^35^.

In this study, we focused on the role of secreted, extracellular AIBP in microglia-driven oxidative stress in AD brain, justifying the choice of a systemic and not microglia-specific *Apoa1bp* knockout mouse. In vitro experiments with microglial cells exposed to Aβ oligomers, to model an AD environment, and treated with rAIBP, to model AIBP gain-of-function, produced results consisted with secreted AIBP loss-of-function in *Apoa1bp^-/-^* APP/PS1 brain. rAIBP significantly reduced Aβ-induced TLR4-inflammarafts, mitochondrial dysfunction and oxidative stress in microglial cells, supporting the hypothesis of a protective role of secreted AIBP in AD brain. However, given the complexity of AIBP biology^8, 24, 36^, we cannot exclude the possibility that intracellular AIBP, both cell-autonomous and internalized, play significant roles in protecting microglia from mitochondrial dysfunction and ER stress. Furthermore, future studies will contribute to the understanding of AIBP effects on neurons, astrocytes and oligodendrocytes.

Another finding of this study supports the relevance of AIBP function to the pathogenesis of human AD, which is more prevalent in women than in men: the AIBP deficiency in female APP/PS1 mice resulted in a more pronounced Aβ plaque accumulation than in males. These findings correlate with the recent report of a significantly altered metabolite profile, including components of cholesterol metabolism, in the blood of female but not male *Apoa1bp^-/-^* mice^37^. AIBP is highly expressed in the mammary gland, ovaries and testes^38^, and secreted AIBP regulates sperm capacitation^39^. This limited set of studies needs to be extended to understand the exact mechanism of sex dependent AIBP effects on cholesterol metabolism in the brain and the development of Alzheimer’s disease.

Taken together, the results of this study demonstrate the expression of TLR4 inflammarafts, which may regulate mitochondrial dysfunction and ER stress in AD microglia. Loss- and gain-of-function experiments suggest that AIBP plays a protective role under conditions of Aβ driven neurodegeneration. Raising levels of a secreted form of AIBP in the brain may be used as a therapeutic strategy to slow down the progression of Alzheimer’s disease.

## METHODS

### Animals

All mouse experiments were conducted in accordance with a protocol approved by the Institutional Animal Care and Use Committee of the University of California, San Diego. Wild type, *Apoa1bp^-/-^* ^14, 15^, APP/PS1^16^, and *Apoa1bp^-/-^* APP/PS1 mice, all on the C57BL/6J background, were housed up to four mice per cage at room temperature, kept in a 12 h light/12 h dark cycle, and received standard laboratory diet and water *ad libitum*.

### Human brain tissue

Human biopsy blocks of brain cortex tissue were obtained from Robert D. Terry as described^40^. Serial sections of 60 nm thickness were sliced from the blocks, placed on silicon wafers, and serial section SEM data was collected on a Zeiss Merlin SEM equipped with OnPoint backscatter detector system at 2.5 kV EHT with 5 nm XY pixels, 12.8 µs dwell time^41^. Each set of images was stacked and aligned using IMOD^42^ to create the volume.

### Mouse tissue preparation

Mice were euthanized using carbon dioxide and perfused with PBS. Brains were removed and dissected into two hemispheres. The hemibrain was post-fixed in 4% paraformaldehyde (PFA) at 4 °C for 5 days. The brains were cut serially at 40 µm in thickness by vibratome (Leica VT1000S) in cold PBS and stored in a cryo-buffer (40% PBS, 30% ethylene glycol, 30% glycerol) at -20 °C before use.

### Immunohistochemistry

Free-floating sections were washed with TBS and permeabilized with 0.2% Triton-X100 in TBS for 20 min under agitation. Sections were incubated in blocking buffer (1% BSA, 5% normal goat serum, and 5% donkey serum in TBS) for 1 h at room temperature. After applying the appropriate antibodies or reagents and completing the staining process, the samples were washed three times with TBS-T, followed by three washes with TBS. Subsequently, TrueBlack® Plus (Biotium, 23014) was applied for 30 seconds, followed by three additional TBS washes. Finally, the samples were mounted using ProLong™ Gold Antifade with DAPI (Invitrogen P36931). For Aβ and microglia staining, sections were incubated with Mouse-on-Mouse IgG Blocking Solution (Invitrogen R37621) at room temperature for 30 min and rinsed three times with TBS-T (0.05 % tween-20). Subsequently, sections were incubated with 82E1 (IBL 10323, 1:100, for Aβ) and IBA1-Fluorochrome (635) (Wako 013-26471, 1:500) overnight at 4 °C, and washed three times with TBS-T and incubated with anti-mouse Alexa Fluor 488 secondary antibody for 2 h at room temperature. For analysis of microglia morphology, tissues were incubated with IBA1 (WAKO 019-19741, 1:500) overnight at 4 °C, washed three times with TBS-T, and incubated with anti-rabbit Alexa Fluor 488 secondary antibody for 2 h at room temperature. Microglia phenotypes were analyzed by using Image J according to a protocol published in ^43^. For detection of apoptosis and loss of neurons, tissue sections were incubated with MAP2 (Abcam ab32454, 1:500) overnight at 4 °C, washed three times with TBS-T, and incubated with anti–rabbit Alexa Fluor 488 antibody for 2 h at room temperature. Tissues were washed three times with TBS-T and were incubated with a TUNEL reaction mixture (In Situ Cell Death Detection Kit, TMR red, Roche 12156792910) for 1 h at 37 °C. For the negative control, sections were incubated with Label solution. For positive control, sections were incubated with DNase I recombinant protein for 3 h at room temperature. Tissues were washed five times with TBS. The percentage of TUNEL positive per cell and the area of MAP2 positive cells were analyzed by Image J. For the measurement of reactive oxygen species, sections were incubated with 10 µM Dihydroethidium (DHE) (Invitrogen, D11347) in TBS at room temperature for 10 min, protected from light. The integrated intensity of DHE positive signal was measured with Image J. Microscopy was conducted using a Leica SP8 super-resolution confocal microscope, operated in a Lightning deconvolution mode.

### Brain tissue extraction and amyloid β_1-42_ ELISA

Soluble and insoluble Aβ fractions were extracted according to ^44^. In brief, after transcardial perfusion with PBS, hemibrain was dissected on ice to isolate mouse hippocampi and cortices, which were frozen at -80 °C until use. For diethylamine (DEA) extraction (non-plaque-associated amyloid beta), 50 mg tissue was transferred into 1 ml solution of 0.2% DEA in 50 mM NaCl, pH 10, with protease and phosphatase inhibitors, vortexed vigorously and mechanically homogenized, followed by centrifugation at 5,000 × g at 4 °C for 5 min to pellet DEA-insoluble material. The supernatant was ultracentrifuged at 80,000 × g for 1 h at 4°C using a type 50 Ti rotor ultracentrifuge (Beckman Coulter). The supernatant, which contains the soluble fraction, was then neutralized by adding 1/10 volume (10%) of 0.5 M Tris HCl, pH 6.8, and gently vortexed. The neutralized samples were divided into aliquots for ELISA and stored at -80°C. The DEA-insoluble pellet was extracted with 0.5 ml RIPA buffer (Cell Signaling 9806S), plus protease phosphatase inhibitors, and sonicated at 4 watts for 1 min on ice. The mixture was then transferred to a new tube and subjected to ultracentrifugation at 80,000 g for 1 h at 4°C. The resulting supernatant represented the RIPA-soluble protein fraction, while the pellet contained the RIPA-insoluble protein fraction. For formic acid (FA) extraction (deposited plaque-associated amyloid beta), the remaining pellets were sonicated in 0.5 ml 70% FA for 7 min on ice. The homogenates were ultracentrifuged at 80,000 × g for 1 h at 4°C. The supernatant was transferred into a new tube and neutralized using a 20x volume of FA neutralization solution, with brief mixing at room temperature. The mixture was ultracentrifuged at 80,000 × g for 1 h at 4°C, and then 50 µl of the supernatant was diluted into 950 µl of FA neutralization solution (1:20) at room temperature. The mixture was briefly mixed, aliquoted, and stored at -80 °C. The DEA and FA fractions extracted from brain tissue were incubated at 37°C for 5 min prior to loading onto ELISA plates. Aβ_1-42_ was measured using a kit from Thermo Fisher Scientific (KHB3442) according to manufacturer’s instructions, using a SpectraMax M2 plate reader (Molecular Devices).

### Aβ preparation

CHO cells stably expressing human APP751 with the Val717Phe mutation (7PA2 cells)^45–47^ were cultured in 10% FBS/DMEM. Conditioned media from confluent 7PA2 cells or control CHO cells cultured in plain DMEM for 16 h was concentrated 10x by using Amicon Ultra Centrifugal Filter with 3kDa cutoff (Millipore UFC500324). The concentration of Aβ_1-42_ was determined by ELISA assay (Thermo Fisher Scientific KHB3442), aliquoted, and stored at -80°C.

### BV-2 cell culture

BV-2 microglia cell line was cultured in DMEM with 5% FBS and 50 μg/ml gentamicin (Omega Scientific) as described previously^5^. Plates were incubated in a 5% CO_2_ atmosphere at 37 °C. BV-2 cells were pretreated with recombinant AIBP (rAIBP)^48^ for 2 h before adding 7PA2-conditioned media to the final concentration of 300 pM Aβ_42_, or control media, for an additional 24 h.

### Lipid rafts and TLR4 staining in BV-2 cells

BV-2 cells were fixed with 4% PFA for 10 minutes and subsequently incubated with Mouse-on-Mouse IgG Blocking Solution (Invitrogen R37621) at room temperature for 30 min. Cells were incubated with recombinant Cholera toxin B (CTxB)-FITC (Sigma Aldrich C1655) for 1 h to stain for lipid rafts and then with an anti-TLR4 antibody (Abcam ab22048) at 4°C overnight, followed by an anti-mouse Alexa Fluor 594 secondary antibody for 2 h at room temperature, and slides were mounted using ProLong™ Gold Antifade Mountant with DAPI (Invitrogen P36931). Image acquisition was performed using a Leica SP8 confocal microscope and image processing using ImageJ Fiji. Colocalization assessment was executed using JACoP plugin, facilitating the calculation of Manders’ coefficients.

### Proximity ligation assay

TLR4-lipid rafts (TLR4-LR) assemblies were assessed using NaveniFlex reagents (Navinci NaveniFlex GM) according to manufacturer’s instructions. Briefly, BV-2 cells were fixed and sequentially incubated with Mouse-on-Mouse IgG Blocking Solution (Invitrogen R37621) and Navenci blocking solution at 37°C for 60 min each in a pre-heated humidified chamber. Unconjugated CTxB (for lipid raft staining, Sigma Aldrich C9903) was incubated at room temperature for 30 min and washed three times with TBS, and then stained with an anti-TLR4 (Abcam ab22048) and anti-CTxB (Sigma 227040) antibodies at 4°C overnight. The samples were incubated with a mixture of Navenibody 1 and 2 at 37°C for 60 min in a preheated humidified chamber, enzyme A in reaction buffer A for 60 min, enzyme B in reaction buffer B for 30 min, and finally, enzyme C in reaction buffer C (Texas red) for a 90 min 37°C. Cells were mounted using ProLong™ Gold Antifade Mountant with DAPI (Invitrogen P36931) and were imaged using a Leica SP8 super-resolution confocal microscope with Lightning deconvolution, and PLA-puncta were counted using the analyze particles function in Image J.

### TLR4 dimerization assay

The TLR4 dimerization assay employed two TLR4 antibodies for flow cytometry: MTS510, which recognizes TLR4/MD2 as a TLR4 monomer but not a dimer, and SA15-21, which binds to cell surface TLR4 regardless of its dimerization status. The percentage of TLR4 dimers was then calculated based on the measurements of MTS510 and SA15-21 in the same cell suspension^5, 49, 50^. For in vitro experiments, BV-2 cells were washed with ice-cold D-PBS, fixed with 4% formaldehyde for 10 minutes, and subsequently washed with D-PBS. Cells were detached using versene, a gentle non-enzymatic cell dissociation reagent, and washed twice with ice-cold FACS buffer, incubated with 2% normal mouse serum containing an anti-CD16/CD32 antibody (FcγR blocker, BD Pharmingen™ 553142) for 30 min on ice, stained with a 1:100 dilution of PE-conjugated MTS510 antibody (ThermoFisher 12-9924-81) and an APC-conjugated SA15-21 antibody (Biolegend 145406), along with a 1:200 dilution of CTxB-FITC (Sigma C1655), for 1 h on ice. Cells were washed and analyzed using a CytoFLEX flow cytometer (Beckman Coulter). For ex vivo experiments, brain tissue was cut into small pieces and processed using Neural Tissue Dissociation kit (P)-papain (Miltenyi Biotec 130-092-628) according to manufacturer’s protocol. Subsequently, the tissue was resuspended in mix A (papain) for 10 minutes, passed through fire-polished pipettes ten times slowly, and incubated with enzyme A for an additional 10 minutes. Cell suspension was cleared using Debris Removal Solution (Miltenyi Biotec 130-109-398), washed with cold D-PBS and resuspended in D-PBS containing 0.5% BSA. Myelin was removed using Myelin Removal Beads II (Miltenyi Biotec 130-096-733), and cells were blocked with 2% normal mouse serum and an anti-CD16/CD32 antibody (FcγR blocker, BD Pharmingen™ 553142) for 30 min on ice, stained with 1:50 anti-CD11b (PerCP-Cy5.5, Biolegend 101228), anti-CD45 (Brilliant Violet 421, Biolegend 103134), 1:100 PE-conjugated MTS510 antibody (ThermoFisher 12-9924-81), an APC-conjugated SA15-21 antibody (Biolegend 145406), along with 1:200 CTxB-FITC (Sigma C1655), and 1:1000 Ghost Dye™ Red 780 Viability Dye (Cell Signaling 18452) for 1 h on ice and protected from light. Cells were then centrifuged, washed and resuspended in a sorting buffer (2% FBS, 1 mM EDTA, DNase and RNase inhibitors in DPBS), followed by filtration through a 70 µm cell strainer before sorting. ArC™ Amine Reactive Compensation Bead Kit (Thermo A10628) and UltraComp eBeads™ Compensation Beads (Thermo 01-2222-42) were used for compensation. Data were analyzed by FlowJo (BD Bioscience).

### Measurement of reactive oxygen species

Intracellular ROS in BV-2 cells was measured by incubating cells with 1 µM cell-permeant 2’,7’-dichlorodihydrofluorescein diacetate (H2DCFDA, Invitrogen, D399) at 37°C for 30 min.

### Sample preparation and multi-modal imaging correlation

Mice were anesthetized with an intraperitoneal injection of ketamine/xylazine and transcardially perfused with about one minute flush of Ringer’s solution containing heparin and xylocaine, followed by approximately 50 ml of 0.1% glutaraldehyde/4% prilled paraformaldehyde in 1x PBS. The brain was dissected and post-fixed with 4% formaldehyde in PBS on ice for 2h and then cut into 100-μm-thick slices. Cortex sections were washed and cryoprotected for 2h, then freeze-thawed by immersion in liquid nitrogen, with further washes in cryoprotectant and PBS solution. For the immunoperoxidase processes^51^, the brain slices were blocked by incubating for 2h in 10% normal goat serum (NGS) (Vector Laboratories), and were incubated overnight with an anti-IBA1 antibody (WAKO 019-19741, 1:100). Next, sections were incubated in the biotinylated goat anti-rabbit secondary antibody and avidin–biotin–peroxidase complex (ABC Elite; Vector Laboratories). They were washed in PBS and reacted with diaminobenzidine (DAB free base, Sigma) with subsequent washes in PBS. Then, brain slices were imaged with a dissecting scope and a transmission light microscope to identify the vasculature and the images were aligned relative to each other in Photoshop (Adobe). DAB-peroxidase labeled tissues were stained for SBEM imaging as previously described. The tissues were stained in succession with 2% reduced osmium tetroxide, 0.05% thiocarbohydrazide, 2% osmium tetroxide, 2% uranyl acetate, and Walton’s lead aspartate solution, with intervening washes using double distilled water. The specimens were dehydrated with 50%, 70%, 90%, 100% EtOH, and dry acetone solutions and then infiltrated and embedded in Durcupan ACM and placed in a 60°C oven for 48h.

The slice was mounted on the end of a small aluminum rod and a low-resolution microCT volume (TBD: pixel size) was collected at 80 kV (Zeiss Versa 510 XRM). The vasculature pattern was used to locate the area of interest in the embedded section. The block was trimmed down to less than 1 mm × 1 mm in size, mounted to a serial block-face EM (SBEM) specimen rivet using conductive silver epoxy (Ted Pella), and left at 60°C overnight. Following trimming of the block using an ultramicrotome, a higher-resolution microCT scan was collected (∼1 mm pixel size) to allow for precise targeting of the SBEM stage to the ROI. An SBEM volume was collected on a Zeiss Gemini 300 SEM equipped with a Gatan 3View and OnPoint backscatter detector system. The SBEM volumes were collected at 2.5 kV EHT with 2.97 nm XY pixels, 50 nm Z steps and 1 µs dwell time. Focal charge compensation with nitrogen gas was used to mitigate charging artefacts. Following data collection, the images were converted to .mrc format and cross-correlation was used for rigid image alignment of the slices using the IMOD image processing package. For volume analysis, we employed the IMOD software package, which is publicly available and specifically designed for visualizing and analyzing three-dimensional EM datasets (http://bio3d.colorado.edu/imod/). Manual segmentation was conducted using drawing tools to trace and outline the structures of interest across consecutive sections of SBEM images.

To create surface models, the ‘imodmesh’ functions were utilized, connecting closed contours by linking defined contours with those directly above and below them. For more precise length measurements, mitochondria were segmented by drawing a series of connected spheres along the length of each mitochondrion using open contour objects^36^. Morphometric data were extracted from the surface models using the ‘imodinfo’ function. The width of the ER lumen was measured using the measurement tool in IMOD, with an unbiased selection of the ER.

### Oxygen Consumption Rate (OCR)

BV-2 cells (2 × 10^4^ per well) were seeded into Seahorse XF24-well plates. At 24 h, cells were pretreated with either BSA or recombinant AIBP (0.2µg/ml) for 2 h, followed by incubation with Aβ (7PA2 conditioned media). OCR was measured using an XF24 analyzer (Agilent). For OCR analysis, after measuring the basal respiration, oligomycin (2 μg/ml, Sigma), an inhibitor of ATP synthesis, carbonyl cyanide 4-(trifluoromethoxy) phenylhydrazone (FCCP; 1 μM, Sigma), the uncoupler, and rotenone (2 μM, Sigma), an inhibitor of mitochondrial complex I, were sequentially added to measure maximal respiration, ATP-linked respiration and spare respiratory capacity.

### Statistical Analyses

Data were represented as Mean±SEM. Data were analyzed with one-way ANOVA with Tukey’s or Dunnett’s multiple comparison test and were applied for normally distributed variables. Log-rank (Mantel-Cox) test was used for mice survival analysis. Differences with a P value of <0.05 were considered statistically significant. All statistical analyses were conducted using GraphPad Prism, version 10.

## DISCLOSURES

### Conflict of interest

Y.I.M., S.-H.C. and W.-K.J. are inventors listed in patent applications related to the topic of this paper. Y.I.M. is scientific co-founder of Raft Pharmaceuticals LLC. The terms of this arrangement have been reviewed and approved by the University of California, San Diego in accordance with its conflict of interest policies. Other authors declare that they have no competing interests.

## ACKNOWLEDGEMENTS

The authors thank Dr. Nicholas Webster (UC San Diego) for generously sharing access to a flow cytometer in his lab and Dr. Goonho Park (Stanford) for providing CHO cells stably expressing human APP751 carrying the V717F mutation.

## FUNDING

This study was supported by NIH grants AG081037 (to Y.I.M., W.-K.J., M.H.E.), HL136275 (to Y.I.M.), EY031697 (to W.-K.J., G.A.P.), AG081004 (to W.-K.J.), EY034116 (to W.-K.J./K.-Y.K./S.-H.C.), NS129684 (to W.-K.J.), NS047101 (to UCSD Microscopy Core), and VA grants I01BX004848 and IBX005224 (to Dr. Nicholas Webster). Shared instrumentation at the National Center for Microscopy and Imaging Research has been supported by NIH grants MH129261, NS120055, NS108934, AG062479, GM138780, AG065549, OD021784, and the NSF grant 2014862.

---

**Figure S1.**
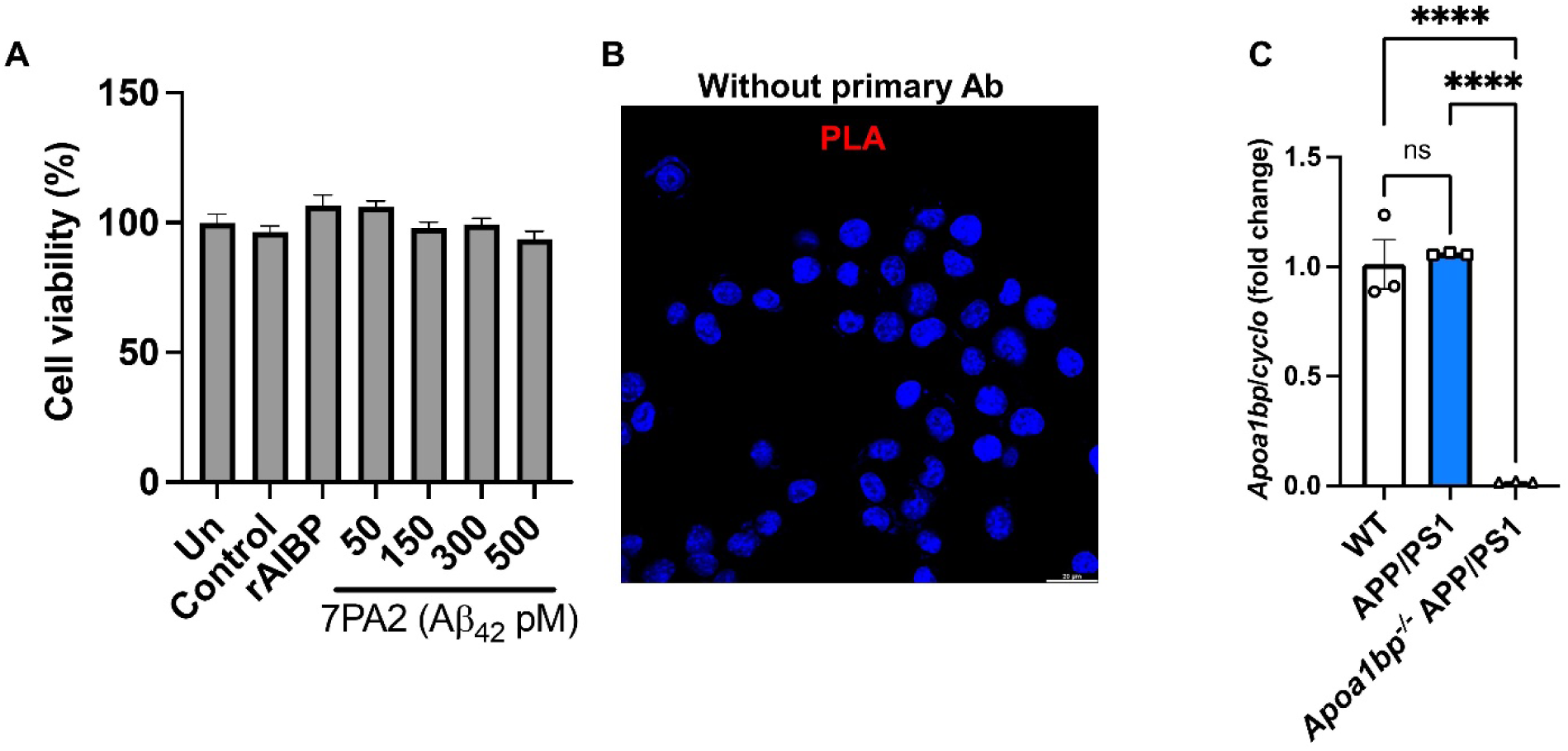
Controls for Figures 1 and 2. (**A**) Cell viability of BV-2 cells incubated with non-conditioned media (Un, unstimulated), control (CHO-conditioned media), 0.5 µg/ml rAIBP, or 7PA2-conditioned media at the indicated concentrations of Aβ for 48 hours (n=7/group). (**B**) In reference to Figures 1C and 1D, image of cells subjected to proximity ligation assay (PLA) in which primary antibodies were omitted. Scale bar, 20 μm. (**C**) Validation of *Apoa1bp* knockout in mouse brain lysates by RT-qPCR: *Apoa1bp* mRNA expression (n=3 per group).

**Figure S2.**
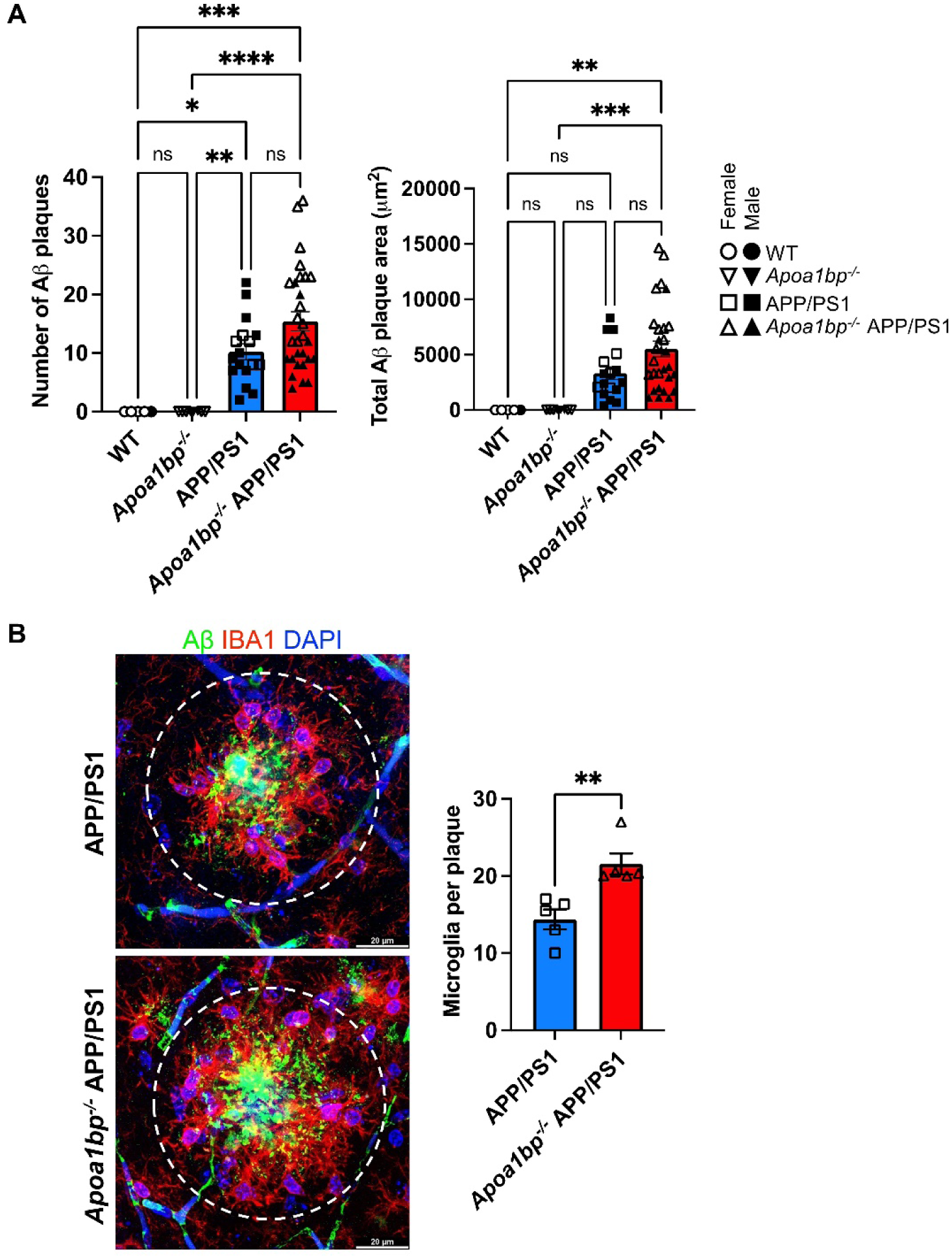
Supplemental data for Figure 7. (**A**) The number of plaques and the total plaque area in the hippocampus of female and male mice (only females are shown in Figure 7). Data are from age-matched 6 months old male and female mice: WT (*n*=5), *Apoa1bp^-/-^* (*n*=8), APP/PS1 (*n*=17), and *Apoa1bp^-/-^* APP/PS1 (*n*=29). Open symbols represent females, and closed symbols represent males. Mean±SEM. One-way ANOVA with Tukey’s multiple comparison test. (**B**) Representative higher magnification image of hippocampus: 82E1 (Aβ; green), IBA1 (microglia; red), and DAPI (nuclei; blue). Scale bar, 20 μm. The numbers of Aβ plaques-associated microglia (inside the circle) were counted in 2-3 different plaque-associated areas in the brain of 5 female mice per group. Mean±SEM. Unpaired t-test.

**Figure S3.**
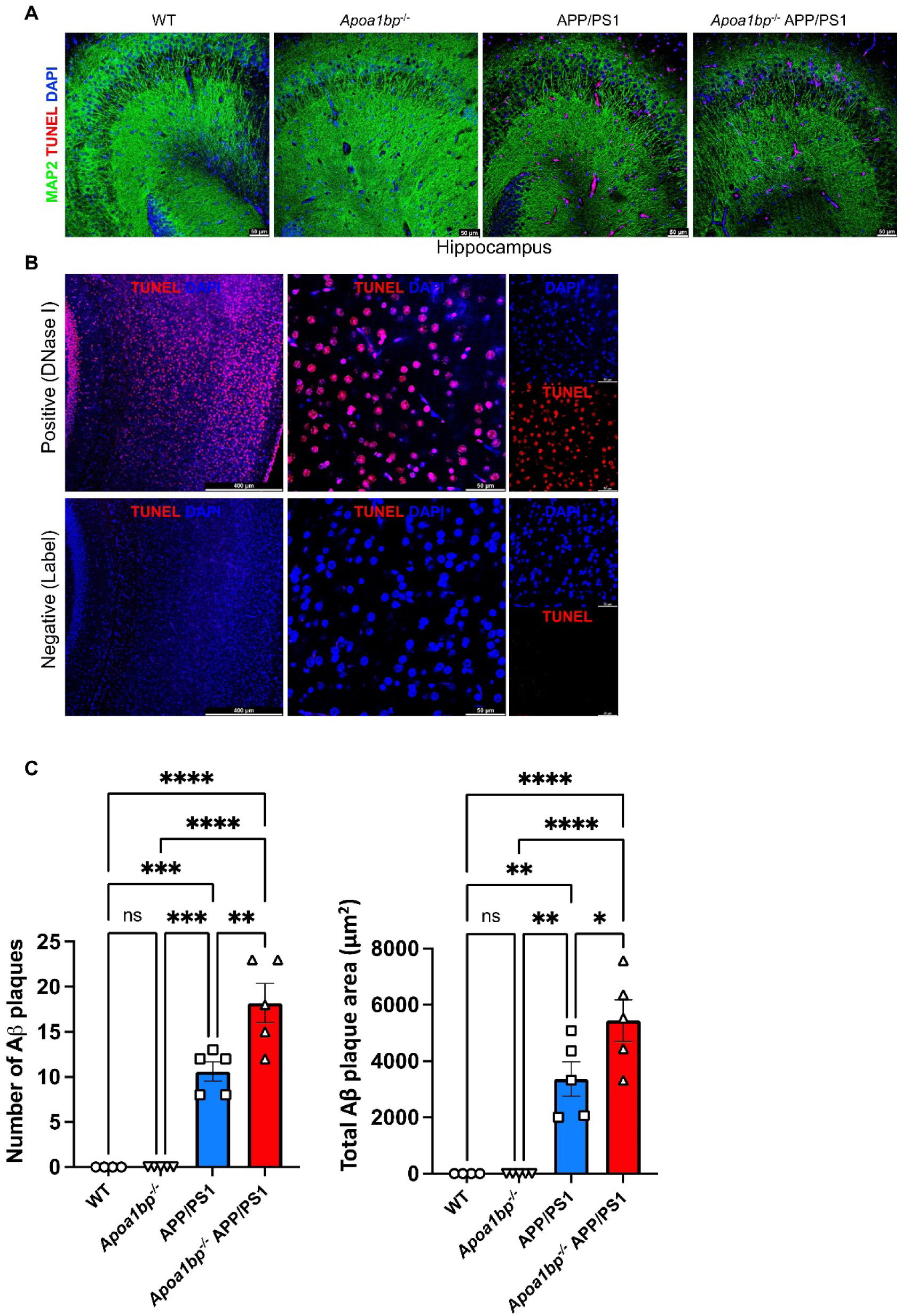
Controls and supplemental data for Figure 8. (**A**) Lower magnification (wider field) images of TUNEL (red), MAP2 (green), and DAPI (blue) staining in hippocampus from female mice. Scale bar, 50 µm. (**B**) Positive control (DNase I) and negative control (Label solution only) of *In Situ* Cell Death Detection TMR red Kit (TUNEL) in female WT mouse brain. Scale bars, 400 µm (left) and 50 µm (center and right). (**C**) The number of plaques and the total plaque area in the hippocampus of mice used for experiment shown in Figure 8 (TUNEL and MAP2). Mean±SEM. One-way ANOVA with Tukey’s multiple comparison test.

## Notes

### Summary of Updates

- changed title - fixed mistake in author's name - small edits in abstract and text

